# Gene coexpression network analysis of galactomannan biosynthesis and endosperm maturation in species of the genus *Coffea*

**DOI:** 10.1101/2024.09.24.614716

**Authors:** Stéphane Dussert, Anna K. Stavrinides, Julien Serret, Virginie Vaissayre, Marie-Christine Combes, Fabienne Morcillo, Eveline Lefort, Stéphanie Rialle, Hervé Etienne, Philippe Lashermes, Thierry Joët

## Abstract

In a few important plant families and genera, including Arecaceae, Fabaceae and the genus *Coffea*, the main seed storage polysaccharide is not starch but cell wall galactomannans. Such seeds are albuminous with a persistent copious living endosperm that accumulates galactomannans. However, our understanding of the regulation of endosperm maturation, cell wall formation and galactomannan biosynthesis in albuminous seeds remains very limited. To gain insights into these processes, a large RNA-seq dataset was produced (14 coffee species × 5 endosperm developmental stages) and scrutinized using gene coexpression network analysis. The network revealed tight transcriptional coordination of the core galactomannan biosynthetic machinery for sucrose import, glycolysis, nucleotide sugar synthesis and transport, arabinogalactan protein and cellulose synthesis, and regulation of the trans-Golgi network. The orchestration of galactomannan and oil accumulation during endosperm maturation appeared to be exerted by the transcription factors FUSCA3, WRINKLED1, SHINE2 and DREB2D. The latter was the only coexpression partner of galactomannan biosynthetic genes. Numerous key genes of galactomannan biosynthesis were significantly upregulated in coffee somatic embryos overexpressing DREB2D, which showed increased production of UDP-galactose and diversion towards raffinose family oligosaccharides. Further, most genes of the galactomannan coexpression module were identified as DREB2D target genes by DAP-seq analysis.

**Highlight:** Gene coexpression network analysis of the maturing endosperm identified the AP2/ERF transcription factor DREB2D as a major regulator of galactomannan accumulation in the cell walls of albuminous coffee seeds.

## INTRODUCTION

Plants store polysaccharides, oil and proteins in seeds to support the emergent young plant after germination until photosynthesis is fully established. Starch is by far the most common polysaccharide stored in seeds (Aguirre *et al*., 2018). In seeds, starch is deposited intracellularly in specialised organelles, the amyloplasts. However, in species of a limited number of genera and families, the main seed polysaccharide reserves are hemicelluloses that accumulate in cell walls, mostly mannans and xyloglucans (Delmer *et al*., 2024), and in even fewer, glucans (Guillon *et al*., 2012). The linear backbone of mannans consists of β-1,4-linked mannose residues with variable degrees of side substitutions. In seeds, the most common form of storage mannans is galactomannans, in which a varying proportion of mannosyl residues is substituted with an α-1,6-linked galactosyl residue (Scheller and Ulvskov, 2010). The degree of galactose substitution affects the solubility of galactomannans in aqueous solutions, their viscosity, and their ability to form a gel (Singh *et al*., 2018; Sharma *et al*., 2022). Thanks to their unique rheological properties, galactomannans are widely used in the food, pharmaceutical, and cosmetic industries (Sharma *et al*., 2022).

Seeds which store mannans in their cell walls are all albuminous seeds with a persistent cellularised living endosperm at the mature stage, thus excluding the caryopsis of cereals, in which the starchy endosperm is a dead tissue. The endosperm of albuminous seeds which store mannans may represent almost the entire seed mass and volume, as is the case in species of the monocot family Arecaceae (palms) and of the dicot genus *Coffea* (Joet *et al*., 2009; Guerin *et al*., 2016). This seed structure is considered plesiomorphic (Forbis *et al*., 2002) and mannan is thought to be the most ancient cell wall hemicellulose (Pauly *et al*., 2013). The family Fabaceae is also known to contain numerous species which produce seeds with a persistent endosperm that accumulates mannans (Buckeridge, 2010). Among these species, seeds of the annual guar (*Cyamopsis tetragonoloba*) or the tree carob (*Ceratonia siliqua*) are exploited for extraction of galactomannan gums that are used in the food industry, mostly as emulsifiers and thickening agents (Singh *et al*., 2018). In these species, the embryo is well developed and the endosperm represents a restrained part of the seed mass, for instance in the carob seed, about half (Laaraj *et al*., 2023).

Over the past three decades, using Fabaceae species that accumulate considerable amounts of galactomannans in their seeds, i.e. guar, fenugreek (*Trigonella foenum-graecum*), senna (*Senna occidentalis*), the genes and enzymes required to synthesize seed storage galactomannans have been identified and characterised. Two enzymes are Golgi-located: the mannan synthase (MANS), which produces the β-1,4-linked mannan backbone (Dhugga *et al*., 2004), and the galactosyltransferase (GMGT), which catalyses the transfer of galactosyl residues from UDP-galactose to the mannan backbone to assemble the galactosyl side chains (Edwards et al., 1999). The post-depositional modulation of the degree of galactose substitution is operated by the cell wall-located α-galactosidase (AGAL)(Edwards *et al*., 1992; Joersbo *et al*., 2001). In addition to these three essential enzymes, another key factor may be needed for seed storage mannan synthesis, the MANNAN-SYNTHESIS RELATED (MSR) protein (Wang *et al*., 2013). In Arabidopsis, AtMSR1 is an auxiliary cofactor that enables the cellulose synthase-like A2 to produce glucomannans instead of mannans (Voiniciuc *et al*., 2019). By contrast, when heterologously expressed in *Pichia pastoris*, coffee MSR1 appeared to be an indispensable cofactor for coffee MANS1 mannan biosynthetic activity (Voiniciuc *et al*., 2019). This finding represents the most recent advance in our understanding of seed storage mannan biosynthesis. However, the transcriptional regulation of this pathway remains largely unknown. More generally, our knowledge of the control of primary wall synthesis is extremely limited (Boerjan *et al*., 2024), in contrast to the considerable progress made in understanding how the synthesis of secondary wall components is regulated through MYB46 activity (Zhong *et al*., 2007). On the one hand, this knowledge gap resembles that concerning the maturation of the endosperm of albuminous seeds, and, on the other hand, that of dicot exalbuminous seeds (Fatihi *et al*., 2016; Alizadeh *et al*., 2021) and of cereal endosperm (Grimault *et al*., 2015 ; Zhang *et al*., 2016b ; Song *et al*., 2024). LEC1-ABI3/FUS3/LEC2 (LAFL) proteins are well-characterised master regulators of embryo maturation in exalbuminous seeds (Fatihi *et al*., 2016; Alizadeh *et al*., 2021). In particular, LEC2 and FUS3 activate the AP2/EREB transcription factor WRINKLED1 (WRI1) (Baud *et al*., 2007; Wang and Perry, 2013), which is thought to play a ubiquitous role in fatty acid synthesis in plants (Ma *et al*., 2013). However, the role of the LAFL regulators and WRI1 in albuminous seeds with a persistent copious endosperm is very poorly documented (Miray *et al*., 2021).

The main reserves in coffee seeds are galactomannans and oil (Joet et al., 2009). Using the cultivation environment as a source of variation in gene expression measured by real time RT-PCR, we previously showed that MANS1, GMGT1 and AGAL1 formed a group of quantitatively coexpressed genes during seed development in *Coffea arabica*, together with two genes coding for enzymes required for galactomannan building block synthesis, mannose-1P guanyltransferase (MGT1) and UDP-glucose 4’-epimerase (UG4E1), which synthesize respectively, GDP-man and UDP-gal (Joet *et al*., 2014). Moreover, time series clustering of microarray data from *C. arabica* seeds at seven developmental stages revealed that MSR1 and a limited number of TF co-clustered with these five genes of the core galactomannan synthetic machinery (Joët *et al*., 2014). Taken together, these two findings suggested that gene coexpression network analysis could help decipher the regulation of endosperm maturation and galactomannan biosynthesis in coffee. Several comprehensive reviews have detailed the advantages of this approach to infer gene function and prioritize candidate genes (e.g. Usadel *et al*., 2009; Li *et al*., 2015). It is worth noting that the two pioneer studies that used coexpression analysis in plant biology enabled the discovery of key players of cellulose and secondary wall biosynthesis in Arabidopsis (Brown *et al*., 2005; Persson *et al*., 2005). Since then, the successful use of gene coexpression network analysis to examine cell wall and oil biosynthesis in plants of major importance in both agriculture and forestry has been reported by numerous authors (e.g. Mochida *et al*., 2011; Guerin *et al*., 2016; Cui *et al*., 2021; Quan *et al*., 2021).

When designing a coexpression study, the challenge is to obtain large but continuous variation in gene expression levels. In the present work, we took advantage of the conserved transcriptional programme of the developing endosperm in fourteen closely related *Coffea* species which differ in the galactomannan and oil contents of their seeds. Using a large RNA-seq dataset (69 transcriptomes), we built a gene coexpression network of the coffee endosperm and searched for novel regulators of galactomannan biosynthesis. Coexpression network analysis identified the AP2/ERF transcription factor DREB2D as a candidate for galactomannan biosynthesis regulation. Its function was then investigated through overexpression in coffee somatic embryos and DNA affinity purification sequencing (DAP-seq) analysis.

## MATERIAL AND METHODS

### Seed material

Seeds from fourteen coffee species conserved in the field genebank of the *Coffea* Biological Resources Center (BRC Coffea, maintained by IRD and CIRAD on Reunion Island) were used for transcriptome and chemical analyses. The fourteen *Coffea* species were *C. arabica* (ARA), *C. brevipes* (BRE), *C. canephora* (CAN), *C.* sp. ‘Congo’ (CON), *C. dewevrei* (DEW), *C. eugenioides* (EUG), *C. heterocalyx* (HET), *C. liberica* (LIB), *C. mauritiana* (MAUR), *C. pocsii* (POC), *C. pseudozanguebariae* (PSEU), *C. salvatrix* (SAL), *C. sessiliflora* (SES), *C. stenophylla* (STE). Seeds at five different maturation stages known to span the development of the endosperm (ST3 to ST7, Joët et al., 2009) were collected from three trees (of three distinct genotypes) per species. As the seed development stage varies considerably among the fourteen species studied, i.e. from ca. 3 to 12 months (Dussert et al., 2000), the developmental stages were based on marked anatomical and morphological seed and fruit traits that are shared across coffee species, as defined and described previously in *C. arabica* (Joët *et al*., 2009; Dussert *et al*., 2018). Briefly, stage 3 coincides with the rapid growth of the aqueous endosperm tissue which progressively replaces the perisperm in the locule (endosperm occupying half of the locule); at stage 4, the remaining perisperm resembles a thin green pellicle surrounding a soft milky endosperm; stage 5 is the peak of reserve deposition and corresponds to endosperm hardening due to massive deposition of galactomannans in the cell walls; stage 6 coincides with fruit veraison, while stage 7 corresponds to mature cherry fruits with red pericarp. After being cross sectioned, the seed was separated from the pericarp and immediately frozen in liquid nitrogen and stored at -80 °C. The endosperm was separated from the perisperm and the embryo while frozen. To minimise the genotypic effect, for each combination of species × development stage, endosperms from the three different accessions collected were evenly pooled prior to grinding and extraction of RNA, lipids, sugars and cell wall polysaccharides (CWP).

### Production of transgenic coffee somatic embryos overexpressing the DREB2D gene

Coffee DREB2D cDNA (Genbank accession number XM_027236031), which was used as a template for PCR amplification, was cloned into the plant overexpression vector pMDC32, transgene expression being driven by a double sequence of the cauliflower mosaic virus 35S promoter. Coffee embryogenic calli (*C. arabica* cv. Caturra) were genetically transformed using recombinant *Agrobacterium tumefaciens* strain LB1119 containing the recombinant plasmid, as previously described (Ribas *et al*., 2011). Somatic embryos regenerated from each hygromycin resistant callus (independent transgenic lines) were tested for transformation by PCR amplification of the selection gene *HPTII*.

### Transcriptome analysis

For each of the 69 combinations of species × development stage (it was impossible to sample seeds at stage 3 in *C.* sp. Congo), endosperms were ground to a fine powder in an analytical grinder (IKA A10, Staufen, Germany) and total RNA was extracted from 70 mg of the powder using the Qiagen RNeasy Lipid Tissue Kit (Qiagen, Stanford, CA). Similar RNA extraction procedures were followed for transgenic somatic embryos. cDNA libraries were constructed using the TruSeq^TM^ Stranded mRNA Sample Preparation Kit (Illumina, USA), then sequenced on an Illumina HiSeq 2500 (single reads, 100 nt) at the MGX platform (Montpellier Genomix, http://www.mgx.cnrs.fr/). After quality filtering using Cutadapt (quality score > Q30 and removal of reads shorter than 60 bp or longer than 140 bp), a minimum of 13 million reads per library were obtained (average of 27.4 million reads per library; Supplementary Table S1). The entire dataset has been deposited at the European Nucleotide Archive (ENA) under project numbers PRJEB32533 and PRJEB79959. Owing to the very low genetic divergence between the *Coffea* species (average of 1.3% gene sequence variation (Cenci *et al*., 2012)), the trimmed reads of each library were mapped to the *C. canephora* coding transcriptome DNA reference sequence (25,574 CDS) (Denoeud *et al*., 2014) using BWA MEM (Li, 2013) with the default parameters, resulting in 60.9-76.3% of mapped reads, independent of the species (Supplementary Table S1). The percentage of mapped reads did not differ significantly among the fourteen coffee species (F_1,13_=1.49; P= 0.149). Reads were counted using IDXstats in SAMtools (Li *et al*., 2009) and read counts were then normalized (RPKM).

Genes with extremely low expression (< 20 counts in total across the 70 tanscriptomes) were not used for subsequent analyses. The R-based DESeq2 software package (Love *et al*., 2014) was used to normalize RNAseq expression values for somatic embryos and to identify genes displaying significantly different expression levels between the controls and DREB2D-overexpressing lines, using a padj cutoff of 0.01. Mapman4 gene ontology annotation (Schwacke *et al*., 2019) was performed using Mercator (Lohse *et al*., 2014). The significance of functional term enrichment in each category of candidates was based on a FDR-adjusted hypergeometric distribution of Mapman BIN categories, using 23,797 genes with annotated BIN categories as a reference.

Transgenic *C. arabica* lines were also tested for overexpression of *DREB2D* and downstream putative target genes using qPCR analysis as described in Joët *et al*. (2014). The set of primers that enabled amplification of target genes is detailed in Supplementary Table S2. The level of expression of each gene was normalized to the geometric mean of expression levels of three validated coffee reference genes (Cc08g05690 Ubiquitin UBQ10, Cc00g15790 40S ribosomal protein S24, and Cc00g17460 14-3-3 protein; Cruz *et al*., 2009).

### Coexpression analysis

Gene coexpression network analysis was performed using the procedure previously described in Guerin *et al*. (2016) after logarithmic transformation of read counts. The strategy used to build the coexpression network of galactomannan biosynthesis during coffee seed maturation combined both the guide-gene approach and the non-targeted approach, as defined by Aoki *et al*. (2007). The network was indeed constructed in the vicinity of a set of galactomannan biosynthetic guide genes, but their partners may belong to the whole coffee genome. The six guide genes used were *MANS1*, *GMGT1*, *MGT1* and *UG4E1-3* genes, all described in a previous transcriptome analysis (Joët *et al*., 2014). These genes encode the enzymes required to produce the nucleotide sugar building blocks GDP-mannose and UDP-galactose (mannose-1P guanyltransferase, MGT, and UDP-glucose 4’-epimerase, UG4E, respectively), to assemble the mannan backbone (mannan synthase, MANS), and to introduce the galactosyl side chains (galactosyltransferase, GMGT). During the first round of network construction, all positive connections between guide genes and their partners (the genes to which they are linked, hereafter termed P1) were computed. During the second round of network construction, partners of P1 were identified (P2) and then links between P2 were generated. Linear regressions were computed using in-house scripts. Two rounds of coexpression analysis using a |R| threshold of 0.85 provided a tractable number of nodes (180) and edges (4878) for network construction. Gene interactions were visualized using the open source software Cytoscape (Shannon *et al*., 2003) and an organic layout. The Markov Cluster (MCL) algorithm (Enright *et al*., 2002) was used for module detection with an inflation of 3.5. Genes of the galactomannan biosynthesis (GMB) coexpression network were manually curated, taking into account recent relevant literature and Mapman annotation using Mercator software to retrieve Mapman functional categories (Lohse *et al*., 2014). Gene names were allocated based on the best Arabidopsis matches.

### Determination of lipid and sugar contents and monosaccharide composition of cell wall material

Total lipids were extracted from 300-mg samples of freeze-dried endosperm powder using a modified Folch method (Laffargue *et al*., 2007). Fatty acid methyl esters (FAMEs) were prepared according to the ISO-5509 standard, and GC analyses of FAMEs were performed as described previously (Laffargue *et al*., 2007). Sugars were extracted and measured by high-performance anion exchange chromatography coupled with pulsed amperometric detection (Dionex Chromatography Co., Sunnyvale, CA, USA) as detailed elsewhere (Dussert *et al*., 2006). Briefly, ca. 50 mg (the exact weight was recorded) of freeze-dried powder were homogenised in 3 ml of 80% (v/v) ethyl alcohol containing lactose (250 mg l^−1^) as internal standard. The mixture was heated for 20 min at 80 °C, during which it was vortexed four times. After cooling in an ice bath, the mixture was centrifuged at 3500 g at 4 °C for 15 min. A 2.5-ml aliquot of the supernatant was collected and dried under reduced pressure in a Speed-Vac concentrator (Jouan RC1010, Saint-Herblain, France). Dried extracts were then dissolved in 9 ml of distilled water and filtered (0.22 μm pore diameter) before analysis. Sugars were quantitated by HPAEC-PAD (Dionex Chromatography Co., Sunnyvale, CA, USA). Sugars were separated on a CarboPac PA-1 (Dionex) column using an isocratic 60 mM NaOH elution at 1 ml min^−1^. Sugar contents were determined by comparison with retention times and calibration curves obtained using sugar standards. The CWP content of the endosperm was measured using the defatted alcohol-insoluble residue (DAIR) method (Redgwell *et al*., 2003). Briefly, about 200 mg (the exact weight was recorded) of freeze-dried powder were homogenized in 5 ml of 80% (v/v) ethyl alcohol. The mixture was heated at 80 °C for 20 min, during which it was vortexed four times. After being cooled in an ice bath, the mixture was centrifuged at 3320 g at 4 °C for 15 min. The supernatant was discarded and the pellet was treated a second time using the same procedure. Lipids were then extracted from the residue twice by homogenizing the pellet in 5 ml of methylene chloride/methanol (2:1), followed by centrifugation at 3320 g for 15 min and removal of the supernatant. After evaporation to dryness under nitrogen at 40 °C, the DAIR was quantified and further treated for monosaccharide composition analysis following the procedure described by Foster et al., (2010). Lipids, sugars and CWPs were all analysed in triplicate (from three different extractions) using a completely random experimental design.

### Recombinant DREB2D protein purification and electrophoretic mobility shift assay (EMSA)

The CDS for *DREB2D* was amplified from cDNA of *C. arabica* seeds using a primer pair containing restriction sites (5‘-ATCA**GGATCC**GAAGCTGACCGTAGTG-3’, and 5’-ATAT**CTCGAG**TCAGTTCCAGGGGTAGTTG-3’). The PCR product was cloned into the bacterial expression vector pET28(a) in frame with the N-terminal 6xHis tag by restriction digest. Recombinant proteins were produced in *Escherichia coli* DE3 (BL21) cells. Transformed cells were grown overnight at 37 °C, with 180 rpm shaking, in LB medium containing kanamycin (30 mg l^-1^). Cultures were then diluted 1:100 in fresh 2 x YT media supplemented with kanamycin, and grown to an OD_600_ of 0.8, before chilling on ice. After induction by 0.1 mM isopropylthio-β-galactoside and overnight culture at 16 °C, bacteria were pelleted by centrifugation and lysed by sonication in Buffer A (50 mM phosphate buffer at pH 8, 50 mM glycine, 500 mM NaCl, 5% glycerol, 10 mM imidazole) supplemented with a protease inhibitor cocktail (Promega) and 0.2 mg ml^-1^ lysozyme. Cellular debris was pelleted by centrifugation at 17000 x g for 15 min at 4 °C, and the supernatant recovered and used to purify polyhistidine-tagged fusion proteins by incubation with nickel-nitrilotriacetic acid agarose resin (Qiagen) for 1 h at 4 °C. Soluble, tagged proteins were purified by washing the resin with Buffer B (the same as Buffer A, with 20mM imidazole), and then eluted in Buffer C (the same as Buffer A, with 100 mM imidazole). Proteins were dialysed in Buffer D (50 mM Hepes buffer pH 7, 100 mM NaCl, 1 % glycerol, 1 mM DTT) with Amicon filtration units (10 KDa cutoff). The concentration of proteins was measured with Bradford reagent (Sigma-Aldrich) according to the manufacturer’s instructions. Proteins were aliquoted, flash-frozen in liquid nitrogen and stored at -20 °C.

Oligonucleotide probes were designed based on the literature to obtain a core DREB recognition site (CCGAC), or with mutation to the DRE element (Feng *et al*., 2019). Probes were synthesized (Thermo Fisher) as single strands (Supplementary Table S3) and were hybridized by mixing the forward and reverse strand in equimolar amounts, heating to 95 °C and then slowly cooling to room temperature. The DRE forward single strand was tagged with AlexaFluor 488 on the 5’ end to serve as labelled probe for the gel shift assays. Test solutions consisted of combinations of DRE fluorescent probe (at 200 or 100 ng per lane) and test probes with or without CcDREB2D. These solutions were mixed and incubated for 25 min before adding loading dye (TBE buffer, 80% glycerol, 0.2% bromophenol blue) and migrating in native gels (10% polyacrylamide, 0.5 X TBE, ran at 100 V). The resulting gels were visualized using a Typhoon FLA 9500 imager (Amersham Biosciences) using the default settings for AlexaFluor488.

### DNA affinity purification sequencing (DAP-seq) and analysis

*Coffea canephora* genomic DNA libraries were prepared for amp-DAP experiments and amplified using the NEBNext Ultra II DNA Library Prep Kit for Illumina (New England BioLabs, UK). The Halo-DREB2D fusion protein was obtained by coupled *in vitro* transcription/translation for 2 h at 27 °C using the TNT SP6 High-Yield Wheat Germ Protein Expression System (Promega, WI, USA). Halo-DREB2D synthesis was verified by Western blot using Anti-HaloTag® Monoclonal Antibodies (Promega). Halo-DREB2D purification and DNA affinity binding were performed following the detailed protocol described by Bartlett *et al*. (2017) and optimized by Hutin *et al*. (2023). Briefly, Halo-DREB2D proteins were immobilised on 20 µl of ChromoTek Halo-Trap Agarose Magnetic Beads for 1 h under agitation on a rotating wheel at 4 °C using 130 µl of cold DAP buffer (PBS supplemented with protease inhibitors (Roche), Nonidet P-40 (0.005% v/v), and 1 mM TCEP (tris(2-carboxyéthyl) phosphine). Agarose beads complexed with Halo-DREB2D were then immobilised on a prechilled magnetic rack, washed 6 times with cold DAP buffer, resuspended in 100 µl of DAP buffer and incubated with 50 ng of gDNA librairies for 90 min at 4°C on a rotating wheel. Beads were washed 10 times with 100 µl of DAP buffer, resuspended in 30 µl of elution buffer (Qiagen) and the bound DNA was eluted from the beads by heating at 98 °C for 10 min. A final DNA amplification step was carried out using the KAPA HiFi Real-Time PCR Library Amplification Kit (Kapa Biosystems, Boston, USA). Amp-DAP DNA libraries were purified with Agencourt AMPure XP beads (Beckman Coulter, Miami, FL) and sequenced on an Illumina NovaSeq 6000 on the MGX platform (Montpellier Genomix). The input gDNA libraries and Halo-DREB2D amp-DAP experiments were produced and sequenced in triplicate.

Read quality was analysed with FastQC (v0.11.9) and screening for contamination was performed using FastQ Screen (v0.15.1). The reads were then aligned with BWA (v0.7.17-r1188) with the -q 30 option on the *Coffea canephora* genome (Denoeud, 2014). The duplicate reads were removed using SAMtools (v1.17). After quality filtering, a minimum of 80 million paired-end reads per library was obtained. Peaks for each replicate were called for separately by comparing their alignment file with that of the control input DNA using MACS2 (v2.7.1) with option fe-cutoff 3 (Zhang *et al*., 2008). Peaks identified on individual replicates were compared using bedtools (v2.31.0) ‘multiinter’ function while the peak annotation was carried out using bedtools “closest” function to assign peaks to genes based on their proximity to the transcription start site (TSS), with a threshold of 2 kb or 4 kb upstream of the TSS. The MEME tool from MEME Suite (v5.5.5) was used to analyse the bound regions to identify enriched DNA sequence motifs, and Tomtom and RSAT tools were used to compare the detected motifs with databases (Bailey *et al*., 2015; Santana-Garcia *et al*., 2022).

## RESULTS

### Distribution of seed lipid and cell-wall polysaccharide contents

The 14 coffee species studied displayed considerable variability in the cell wall polysaccharide (CWP) content of their seeds (Fig. 1A), which ranged from 47.5% dry matter (DM) in *C. salvatrix* to 81.1% DM in *C. mauritiana*. The total lipid content of mature seeds also varied considerably from 10.8% DM in *C. mauritiana* to 29.6% DM in *C. pseudozanguebariae* (Fig. 1B). By contrast, the fatty acid composition of total lipids was very similar among the fourteen species, palmitic and linoleic acids each accounted for 35-45% of total fatty acids (Supplementary Table S4). The cell wall polysaccharides (CWP) and lipid contents were highly negatively correlated (R^2^=0.79, Fig. 1C), indicating that CWP and lipids collectively represent most of the mature endosperm reserves in all coffee species.

**Figure 1.**
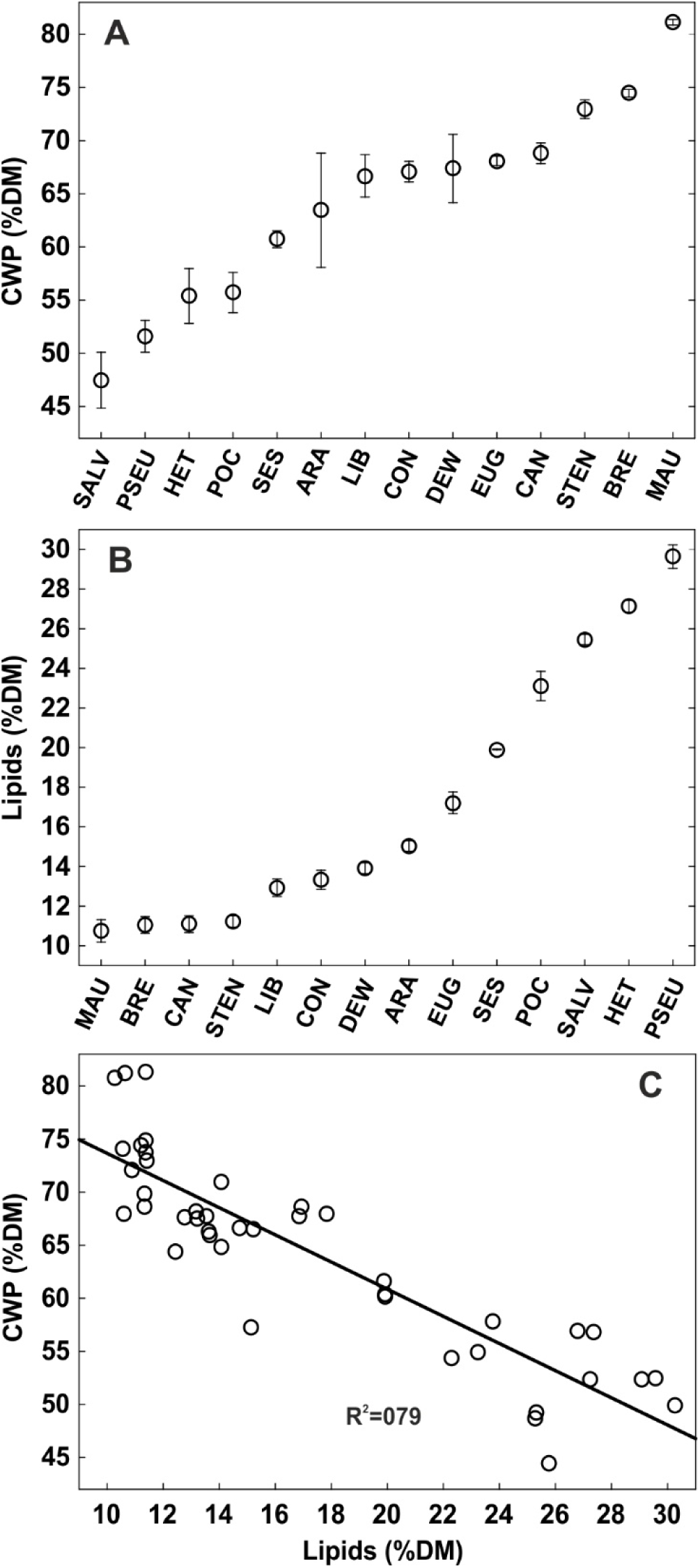
Distribution of cell wall polysaccharides and lipid contents (% DM) in mature seeds of 14 *Coffea* species. Distribution of seed cell wall polysaccharide (A) and oil (B) contents in the 14 *Coffea* species studied. (C) Negative correlation between cell wall polysaccharide and oil contents. The 14 *Coffea* species are *C. arabica* (ARA), *C. brevipes* (BRE), *C. canephora* (CAN), *C.* sp. Congo (CON), *C. dewevrei* (DEW), *C. eugenioides* (EUG), *C. heterocalyx* (HET), *C. liberica* (LIB), *C. mauritiana* (MAUR), *C. pocsii* (POC), *C. pseudozanguebariae* (PSEU), *C. salvatrix* (SAL), *C. sessiliflora* (SES), *C. stenophylla* (STE).

### The transcriptional programme of endosperm development and maturation is conserved between *Coffea* species

The conservation of the transcriptional programme of the developing endosperm in the fourteen species was studied using genes known to be expressed at specific developmental stages in *C. arabica* (Joët *et al.,* 2009; Dussert *et al*., 2018; Combes *et al*., 2022) (Fig. 2). First, in all species (except *C. mauritiana* and *C. pseudozanguebariae*, which do not produce caffeine) the key genes for caffeine biosynthesis (namely *XMT*, *MXMT* and *DXMT*) were highly expressed at early stages 3 and 4, when the endosperm develops rapidly and replaces the perisperm in the locule, followed by a sharp decrease in transcript levels at later stages (Fig. 2A). Second, transcripts for key genes of galactomannan metabolism (MANS1, GMGT1, AGAL1) exhibited a highly conserved accumulation pattern in all fourteen coffee species, *i.e.* a transient and pronounced peak in mRNA observed at stages 4 and 5 (Fig. 2B). Stages 4 and 5 coincide with the storage phase in *C. arabica*, including endosperm expansion at stage 4 and endosperm hardening due to the massive deposition of galactomannans in cell walls at stage 5 (Joët *et al*., 2009). Third, transcript profiling of genes which act specifically during late maturation and acquisition of desiccation tolerance, *i.e.* genes coding for the dehydrin DH1, the heat shock protein HSP90, and the BURP domain dehydration-responsive protein RD22-like (Stavrinides *et al*., 2020), showed that this developmental process takes place at stages 6 and 7 in all *Coffea* species and is distinct from the storage stage (Fig. 2C). Finally, both hierarchical cluster analysis (HCA) and principal component analysis (PCA) of the whole transcriptome dataset showed grouping of samples at stages 3 and 4 in almost all species, grouping of samples of stages 6 and 7 in a second group, with stage 5 endosperms between the two groups (Supplementary Fig. S1). Taken together, these results indicate a conserved transcriptional sequence of endosperm development in the genus *Coffea*, with major distinct switches governing endosperm growth, reserve deposition, and late maturation, independent of the total duration of the seed development stage, which varies from 3 to 12 months in the fourteen coffee species studied (Dussert *et al*., 2000).

**Figure 2.**
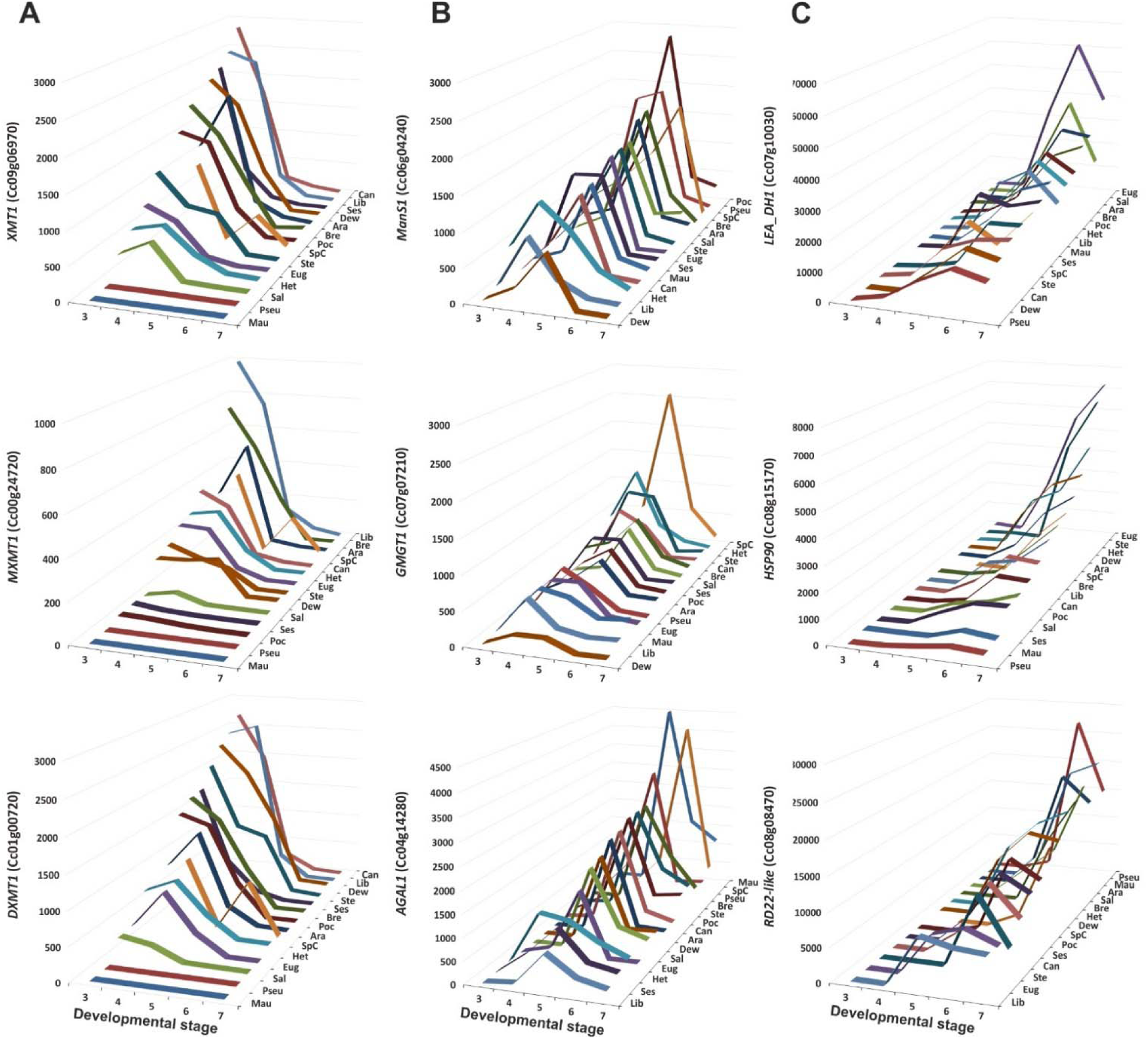
Expression profiles (rpkm) across the development of stage-specific genes in the seeds of the 14 *Coffea* species studied. **A.** Early endosperm development: genes involved in caffeine biosynthesis. *DXMT1* (Cc01g00720), 3,7-dimethylxanthine methyltransferase (caffeine synthase); *MXMT1* (Cc00g24720), 7-methylxanthine methyltransferase (theobromine synthase); *XMT1* (Cc09g06970), xanthosine methyltransferase. **B.** Storage phase: genes involved in galactomannan biosynthesis. *AGAL1*, alpha-galactosidase (Cc04g14280), *GMGT1*, galactomannan galactosyl transferase (Cc07g07210), *MANS1*, mannan synthase (Cc06g04240). **C.** Late maturation: genes encoding the heat shock protein HSP90 (Cc08g15170), the dehydrin DH1 (Cc07g10030) and the dehydration-responsive protein RD22-like (Cc08g08470).

### The galactomannan biosynthesis guided gene coexpression network highlights the key processes associated with the synthesis and deposition of cell wall storage polysaccharides

Two rounds of guided coexpression analysis using six key genes of galactomannan biosynthesis as guide genes (MANS1, GMGT1, MGT1 and UG4E1-3), the complete RNA-seq dataset (stages 3 to 7) and a |R| threshold of 0.85, yielded two distinct networks of very contrasted sizes (Fig. 3A). The three *UG4E* guide genes formed a very small network of six genes, while *MANS1*, *GMGT1* and *MGT1* formed a second large network of 174 genes, hereafter referred to as the galactomannan biosynthesis (GMB) network. The Markov clustering algorithm identified three modules in the GMB network. Modules 1, 2 and 3 contained 121, 43 and 11 genes, respectively. The three guide genes (MANS1, MGT1, and GMGT1) all belong to the second module. Module 2 also includes MSR1 (Cc00g16710), a key cofactor recently reported to be indispensable for coffee MANS1 mannan synthase activity (Voiniciuc *et al*., 2019) (Fig. 4).

**Figure 3.**
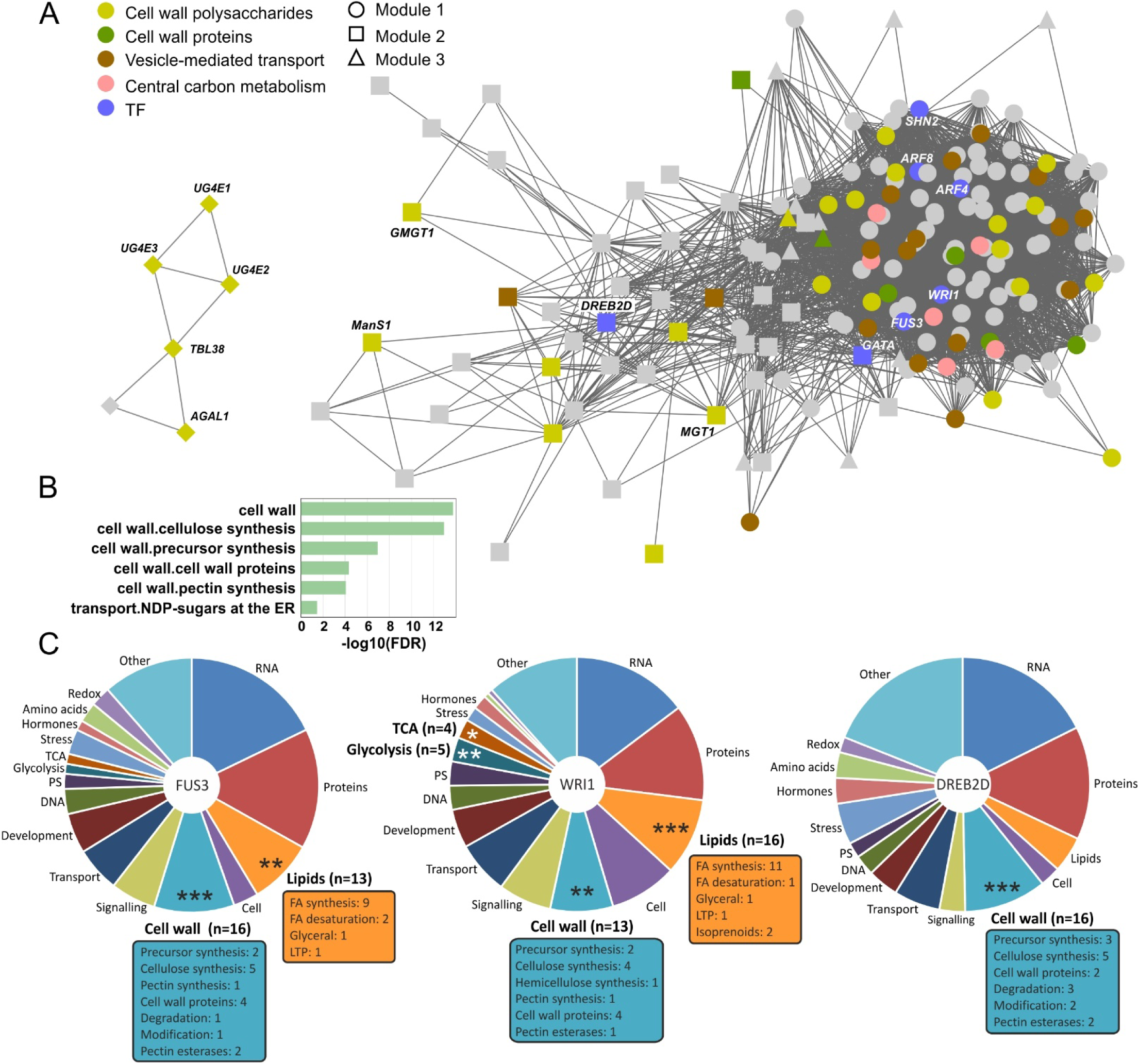
Gene coexpression analysis of galactomannan biosynthesis. **A.** Gene coexpression analysis was performed using the whole transcriptome dataset, six galactomannan-related guide genes (*AGAL1*, *GMGT1*, *MANS1*, *MGT1*, *UG4E1*, *UG4E2*, *UG4E3*), and an |R| threshold of 0.85. The three *UG4E* guide genes formed a very small network of six genes (on the left), while *MANS1*, *GMGT1* and *MGT1* formed a large network comprising 174 genes, the galactomannan biosynthesis (GMB) network (on the right). The three modules detected in the GMB network using the Markov clustering algorithm are represented by different symbols (modules 1, 2 and 3 are represented by respectively, circles, squares and triangles). The most relevant functional groups of the GMB network genes are shown in colour: cell wall polysaccharide and protein synthesis, vesicle-mediated transport, central carbon metabolism, and transcription factors. **B**. False discovery rate (FDR) of the functional terms that showed significant enrichment in genes of the GMB network. **C.** Pie charts of functions assigned to the top 200 coexpression partners of FUS3, WRI1 and DREB2D. Significantly enriched functions are indicated by bold stars and the number of genes involved is given in brackets next to the name of the functional category. The number of genes in the different functional subclasses is given in the coloured boxes. FA, fatty acid biosynthetic genes; PS, photosynthesis-related genes; CWP, cell wall polysaccharides.

**Figure 4.**
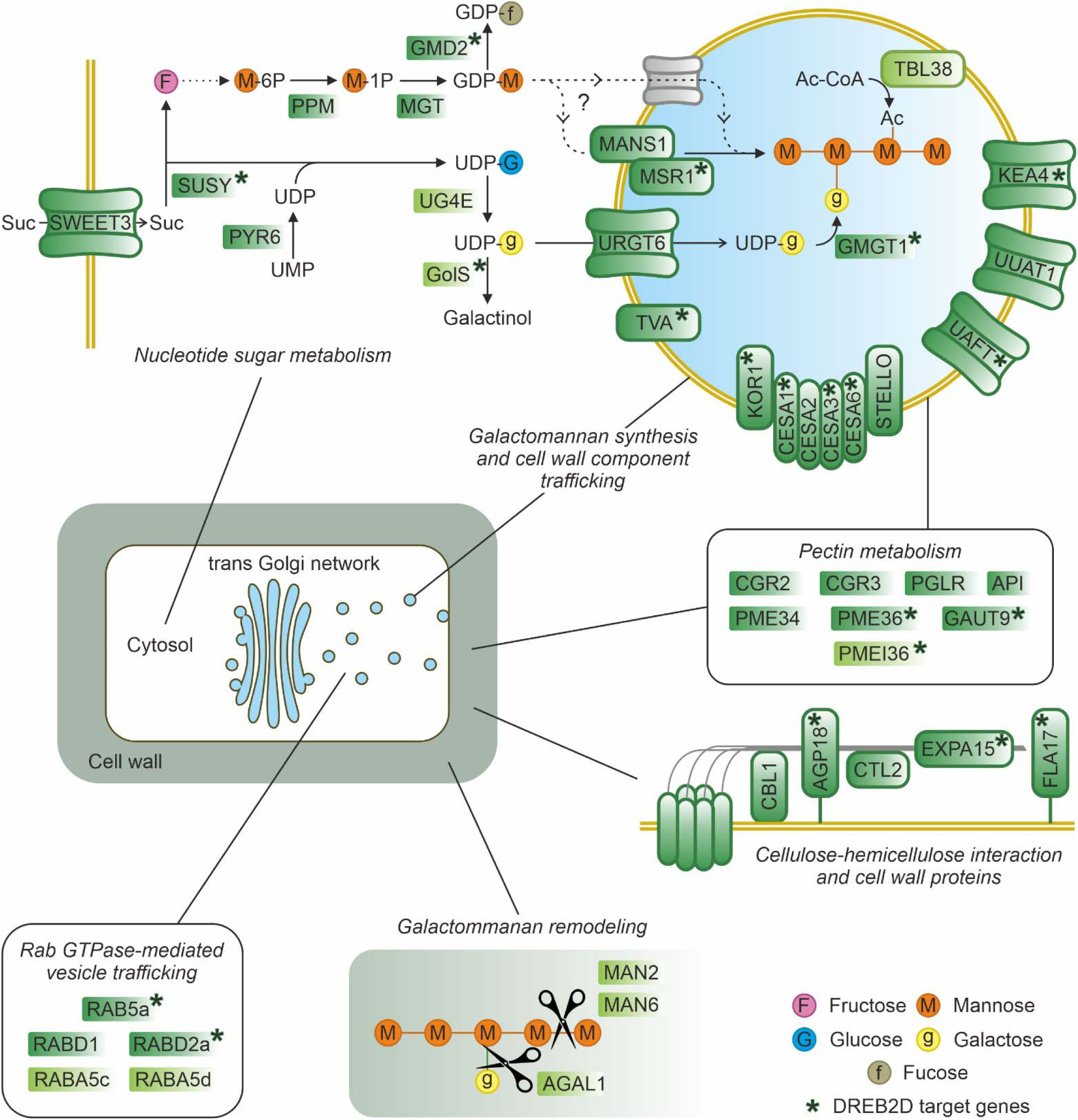
Enzymes and proteins of the GMB coexpression network involved in the synthesis, deposition and modification of cell wall storage polysaccharides in the coffee endosperm. Scheme of the major biosynthetic pathways and metabolic processes occurring in different endosperm cell compartments. All enzyme and protein names are explained in Supplementary Tables S2 and S3. Enzymes and proteins highlighted in dark green are nodes of the GMB coexpression network. Enzymes and proteins followed by an asterisk are encoded by putative DREB2D target genes, as revealed by DAP-seq analysis. Enzymes and proteins highlighted in light green are nodes of the ST3-ST5 gene coexpression network. The scheme was adapted from Figure 2 of the review by Voiniciuc (2022).

Among the 174 genes of the GMB network, 124 genes (71%) were assigned to functional groups and subgroups based on high amino acid sequence homology with proteins that have already been functionally characterised in model plants (Fig. 3B, Supplementary Table S5). The GMB network showed highly significant enrichment (FDR<0.0001) in the functional terms cell wall, cell wall precursor synthesis, cellulose synthesis, pectin synthesis and cell wall proteins (Mapman bins 10, 10.1, 10.2, 10.4 and 10.5, respectively). In all, the cell wall-related genes represented 29% of the coexpression network annotated genes (36 genes). Among them, the coexpression network contains four genes that encode components of the cellulose synthase complex (CSC), i.e. CESA1, CESA2, CESA3, CESA6, and three key proteins of CSC secretory vesicles, KOR1, TRANVIA, and STELLO (Zhang *et al*., 2016) (Fig. 4). TRANVIA (TVA) was recently described as a component of secretory compartments which promotes the delivery of cellulose synthase complexes to the plasma membrane through the trans-Golgi network (Vellosillo *et al*., 2021).

The network also includes COBRA-like protein 1 (CBL1) and Chitinase-like protein 2 (CTL2), both of which are involved in interactions between cellulose microfibrils and hemicellulose (Sánchez-Rodríguez et al., 2012; Aniento et al., 2022) (Fig. 4). Moreover, the presence of seven genes involved in the synthesis of cell wall proteins such as arabinogalactan proteins (AGPs), is also worth noting, as well as that of seven genes encoding well-characterised enzymes devoted to pectin synthesis and modification (CGR2, CGR3, PGLR, PME34, PME36, API, GAUT9) (Fig. 4). Finally, the GMB network also contains six genes dedicated to the biosynthesis and transport of nucleotide sugars (GMD2, PPM, RGP1, Uaft, URGT6, UUAT1), including key players of the GDP-Man pathway, such as the phosphomannomutase (PPM), which catalyses the interconversion of man-6P to man-1P and is essential for the biosynthesis of GDP-man, or URGT6, which transports UDP-gal into the Golgi lumen (Rautengarten *et al*., 2014) (Fig. 4).

Another functional class overrepresented in the GMB network is cellular transport (17 genes), with numerous genes involved in vesicle-mediated transport via the trans-Golgi-network, including KEA4, a Golgi-located cation/proton exchanger which suports Golgi function in cell-wall biosynthesis (Wang *et al*., 2019), SCAMP4, and several RAB-GTPases which play a major role in the regulation of membrane tethering and fusion in the trans-Golgi network and in the secretory pathway of cell wall polysaccharides (Lunn *et al*., 2013) (Fig. 4). Finally, key players in central carbon metabolism are also present in the GMB network, including SWEET3, CIF1, SUSY1, which are involved in sucrose import and cleavage, as well as the glycolytic enzymes HXK-5 and PGLYM-p (Fig. 4).

Together with AGAL1 and Trichome birefringence-like TBL38 (Cc02g14650), which is involved in O-acetylation of cell wall polymers during their deposition in Arabidopsis (Sun *et al*., 2020), the three UG4E genes formed a small network of six genes that was not connected to the main GMB network (Fig. 3A). However, together with TBL38 and AGAL1, UG4E genes joined the GMB network when the ST3-ST5 RNA-seq dataset (42 transcriptomes) was used to build the network (Supplementary Fig. S2; Supplementary Table S6). Developmental stages ST3 to ST5 corresponded to the period of cell wall galactomannan deposition and oil biosynthesis (Joët *et al*., 2009, 2014). This ST3-ST5 network was shown to contain all the guide genes used for coexpression analysis, i.e. *MANS1*, *GMGT1*, *MGT1, UG4E1, UG4E2, UG4E3*, key players of CWP synthesis, such as TVA and MSR1, other enzymes which catalyse mannan backbone modifications, such as the mannan endo-1,4-beta-mannosidases MAN2 and MAN6 (Cc05g07220 and Cc06g00320, respectively), and the galactinol synthase GolS1b (Cc02g35340) involved in the conversion of UDP-galactose into raffinose family oligosaccharides (Fig. 4).

### Gene coexpression analysis identified transcriptional regulators of galactomannan biosynthesis

The GMB network includes seven transcription factors (TFs), namely, three TFs of the B3 superfamily (FUS3 and auxin response factors ARF4 and ARF8), three TFs of the AP2/ERF superfamily (including WRI1, SHN2 and DREB2D) and one GATA TF (Fig. 3A; Supplementary Table S5). Since FUS3 and WRI1 are well-characterised master regulators of seed maturation and oil accumulation in model plants, we investigated the functions of their respective top 200 most positively connected genes according to the value of the Pearson coefficient in coexpression analysis. The top 200 coexpression partners of both FUS3 and WRI1 displayed a significant enrichment for lipid and cell wall related functions, including fatty acid synthesis, cellulose synthesis and cell wall proteins (Fig. 3C). Genes involved in glycolysis and tricarboxylic acid (TCA) cycle were also significantly overrepresented among the top 200 partners of WRI1. Although significantly expressed during endosperm development (Supplementary Fig. S3), the two seed master regulators LEL1 (LEC1-like) and ABI3 were not present in the GMB coexpression network. The expression profile of LEL1 was very similar in all the coffee species, but with a transcription peak at ST3 prior to that in CWP and oil biosynthesis genes. ABI3 did not display a shared uniform transcriptional profile among the fourteen coffee species studied and therefore had no coexpression partners, even at an |R| threshold as low as 0.7 (Supplementary Fig. S3; Supplementary Table S7).

DREB2D (Cc10g02270) belongs to Module 2 of the GMB network and was the only TF that was a direct partner (first round of coexpression analysis) of the guide genes MANS1, GMGT1 and MGT1 (Fig. 3A). It is worth mentioning that TVA and MSR1 were also direct coexpression partners of DREB2D in Module 2, with a Pearson coefficient of 0.90 and 0.93, respectively. Moreover, genes involved in the metabolism of cell wall polysaccharides and cellulose synthesis were significantly enriched in the top 200 coexpression partners of DREB2D (Fig. 3C). Finally, DREB2D was the only TF primarily connected to GMB guide genes, MSR1, TBL38, and MAN6 in the coexpression network built using ST3-ST5 RNA-seq data (Supplementary Fig. S2). Gene coexpression network analysis using galactomannan biosynthetic genes as guide genes therefore provides evidence that DREB2D is a major candidate for direct transcriptional regulation of CWP biosynthesis.

### Overexpression of DREB2D in *C. arabica* somatic embryos altered soluble sugar composition

Transgenic *C. arabica* somatic embryos overexpressing the DREB2D cDNA sequence under the control of the constitutive 35S promoter of the Cauliflower mosaic virus were regenerated using *Agrobacterium*-mediated transformation. Real-time RT-PCR analysis showed a 6-to 250-fold increase in the expression of the DREB2D transgene in the eight independent transformed lines tested compared to control lines (Fig. 5A; Supplementary Table S8). The CWP content of DREB2D+ somatic embryos and controls did not differ significantly (Fig. 5B). No significant differences in the monosaccharide composition of CWPs was observed between control and DREB2D+ lines (Fig. 5C), either for mannose content or the major residues, namely arabinose, glucose, galactose and xylose, which together account for more than 80% of total CWP monosaccharides. By contrast, soluble sugar profiling revealed that DREB2D overexpression altered the soluble sugar content, with a significant increase in raffinose and stachyose contents and a significant decrease in the inositol content (Fig. 5D). Raffinose increased dramatically, with accumulation levels more than 20 times higher in DREB2D+ lines (1.2 ± 0.5 %DM) than in controls (<0.1 %DM).

**Figure 5.**
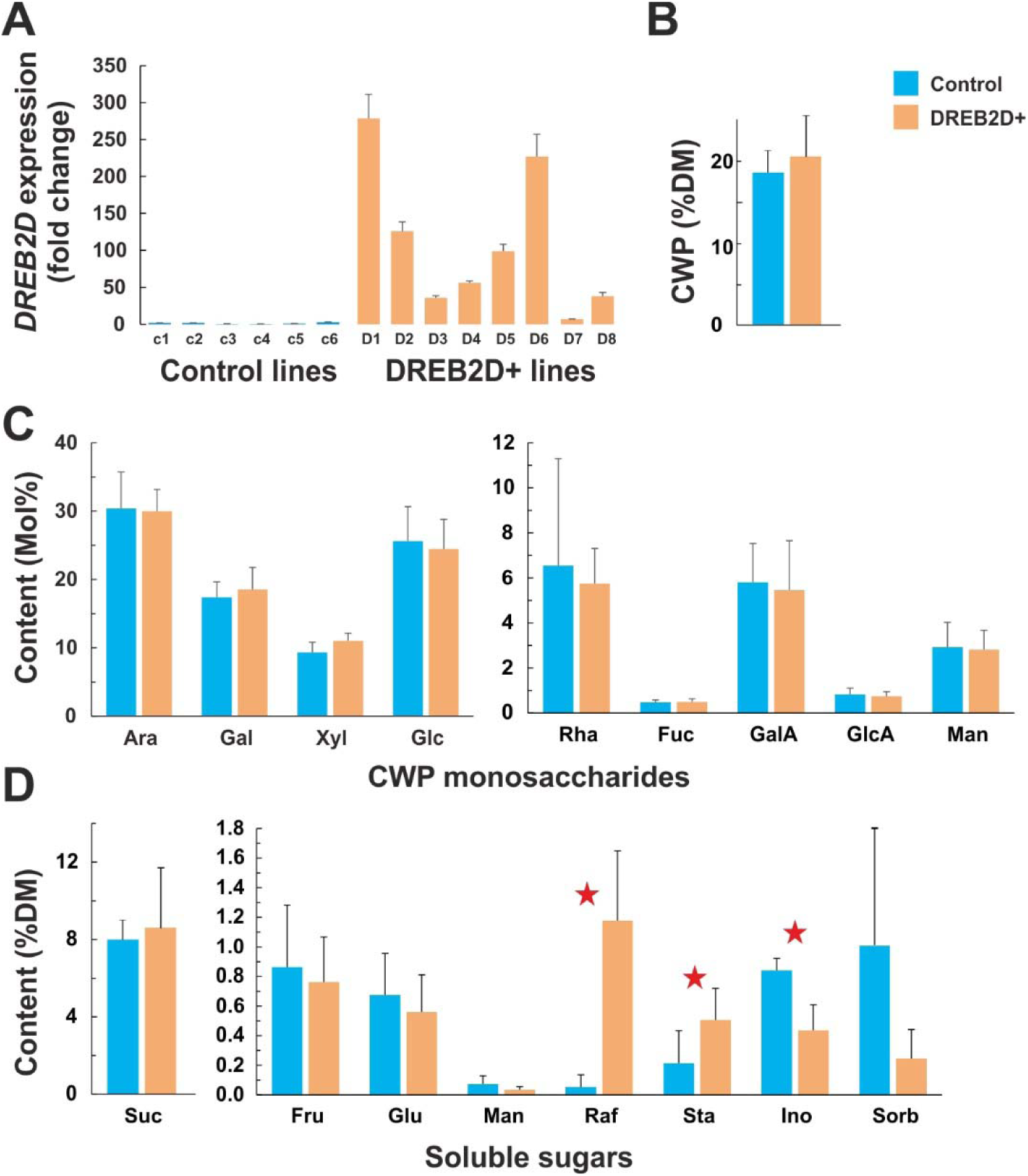
Influence of DREB2D overexpression on the cell wall polysaccharide content and composition and soluble sugar contents of *C. arabica* somatic embryos. **A.** DREBD2 expression level as measured by Real-time RT-PCR in six control and eight DREBD2+ lines. **B.** Cell wall polysaccharide (CWP) content in control and DREBD2+ somatic embryos. **C.** Monosaccharide composition of CWP in control and DREBD2+ somatic embryos. **D.** Soluble sugar contents in control and DREBD2+ somatic embryos. Ara, arabinose; Fru, fructose; Fuc, fucose; Gal, galactose; GlcA, glucuronic acid; GalA, galacturonic acid; Glu, glucose; Ino, myo-inositol; Man, mannose; Raf, raffinose; Rha, rhamnose; Sorb, sorbitol; Sta, stachyose; Suc, sucrose; Xyl, xylose. The effect of DREB2D overexpression was tested using one-way ANOVA. Significant effects are indicated with an asterisk for P < 0.01.

### DREB2D modulates the expression of genes related to cell wall metabolism

An Illumina-based RNAseq experiment was then conducted using six DREB2D+ lines and four control lines (Supplementary Tables S9-S10). Deseq2 analysis revealed 881 differentially expressed genes (DEGs, Padj<0.01), including 532 up-regulated and 349 down-regulated genes in DREB2D+ somatic embryos (Supplementary Tables S11). Among the significantly upregulated genes were several key genes of galactomannan biosynthesis and modification, including those encoding UG4E1, UG4E2, UG4E3, AGAL1, MSR1, TRANVIA, PPM, TBL38, TBL42 and three mannan endo-1,4-beta-mannosidases (Fig. 6A; Supplementary Table S11). Of the 532 up-regulated genes, 110 genes showed a more than 4 times transcriptional induction in DREB2D+ transgenic lines compared to control embryos. Mapman enrichment analysis showed that this top 110 gene subset is significantly enriched in cell wall metabolism and sugar metabolism functions (Fig. 6B). In addition, several late maturation functions were significantly enriched, including ABA metabolism, late embryogenesis abundant (LEA) proteins and abiotic stress response (Supplementary Table S12). In particular, nine genes encoding LEA proteins, as well as nine genes encoding small heat shock proteins (sHSP) were detected among up-regulated DEG in DREB2D+ somatic embryos (Supplementary Tables S11).

**Figure 6.**
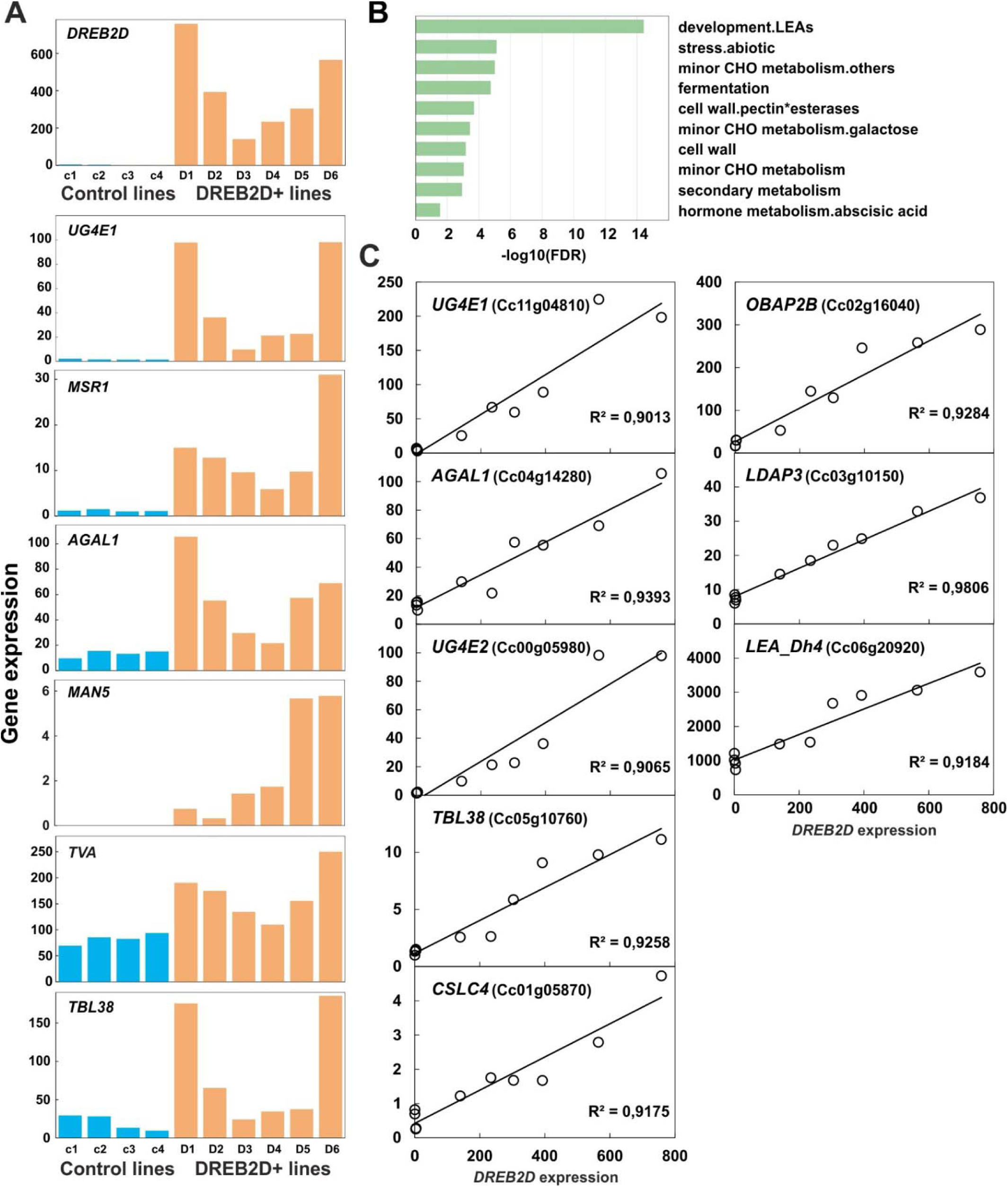
Functional validation of DREB2D in coffee somatic embryos. **A**. Transcript amounts measured by RNA-seq of *DREB2D* and six putative target genes (*UG4E1*, *MSR1*, *AGAL1*, *MAN5*, *TVA* and *TBL38*) in four control lines (blue bars) and six DREBD2+ lines (orange bars). **B.** False discovery rate (FDR) of the functional terms that show significant enrichment in the 110 genes which were massively (4 fold) upregulated in DREB2D+ transgenic somatic embryos compared to in control somatic embryos. Values are means (±SD) of four technical replicates for eight independent and six control lines transformed with empty PMDC vector. Real-time RT-PCR analysis of expression of key galactomannan biosynthetic genes in DREB2D+ transgenic somatic embryos. **C**. Highly significant positive correlations (R > 0.9) between transcript amounts (rpkm) of *DREB2D* and eight putative target genes (*UG4E1, AGAL1, UG4E2, TBL38*, *CSLC4*, *OBAP2D*, *LDAP3*, and *LEA_Dh4*). AGAL1, alpha-galactosidase; AGL, agamous-like; CSLC, cellulose-synthase-like-C; LDAP, lipid-droplet associated protein; LEA-dh, late embryogenesis abundant protein, dehydrin; OBAP2B, oil body associated protein 2B; TBL38, trichome-birefrengence-like, UG4E, UDP-glucose 4-epimerase.

Putative DREB2D target genes were also scrutinized using linear regression, based on the assumption of a quantitative relationship between the expression level of DREB2D and that of genes downstream of DREB2D. Correlations found between DREB2D and target gene expression levels revealed 38 candidate genes with a R^2^ higher than 0.9 (FDR<0.01) (Supplementary Tables S13). This subset of 38 candidate target genes contains key genes of CWP synthesis and modification, including UG4E1, UG4E2, TBL38, AGAL1, and the cellulose synthase-like *CSLC4* (Cc01g05870), whose orthologue is involved in xyloglucan synthesis in Arabidopsis (Kim *et al*., 2020) (Fig. 6C). Finally, the subset of genes quantitatively co-expressed with *DREB2D* in transformed somatic embryos include proteins typical of the seed maturation programme such as the LEA dehydrin Dh4 (Cc06g20920), the oil-body associated protein OBAP2B (Cc02g16040) and the lipid-droplet associated protein LDAP3 (Cc03g10150) (Fig. 6D).

Interpretation of functional enrichment of genes down-regulated in DREB2D+ lines is less straightforward because the cellular context of somatic embryos is very different from that of the endosperm, and these genes may not play any role during seed development. However, alpha-mannosidase genes (Cc04g05900, Cc09g07970), as well as several genes involved in pectin modifications such as polygalacturonases (Cc03g13740, Cc09g05370) and rhamnogalacturonate lyase (Cc10g08340), were significantly down-regulated in somatic embryos that overexpressed DREB2D. Furthermore, Mapman-based enrichment analysis significantly highlighted many processes that co-occur in plastids, including synthesis of thiamine, salicylic acid precursors, monoterpenes and carotenoids (Supplementary Table S12). In addition, many genes encoding components of the photosynthetic electron transfer chain were also detected among down-regulated genes. It is worth noting that visible differences in pigmentation were observed between DREB2D+ and control somatic embryos, most probably related to lower carotenoid and chlorophyll contents in DREB2D+ lines (Supplementary Fig. S4). These results suggest a role for DREB2D in the structural and functional changes in plastids associated with seed maturation. Another striking observation was the presence of all the key genes for caffeine biosynthesis (XMT, MXMT and DXMT) in the set of genes down-regulated in DREB2D+ lines. Since these genes are well-known markers of perisperm and early endosperm developmental stages (Denoeud *et al*., 2014), this result suggests that DREB2D may also act as a broad repressor of key processes of early endosperm development, thereby facilitating and synchronising entry into the maturation phase.

### DREB2D binds the promoter of genes of the GMB coexpression network

Phylogenetic analysis of CcDREB2D together with *Arabidopsis thaliana* members of DREB subfamily A-2 of the APETALA2/ethylene-responsive factor (AP2/ERF) family identified CcDREB2D as a close relative to AtDREB2D (AT1G75490) and AtDREB2G (AT5G18450) (Supplementary Fig. S5). DREB2D belongs to a class of TF proteins known to bind to C-repeat or dehydration response elements (CRT/DRE) in gene promoters (Jiang Chao *et al*., 1996). An electrophoretic mobility shift assay (EMSA) was therefore performed to test CcDREB2D binding to artificial DRE elements *in vitro* (Fig. 7A). Binding of the DREB2D-His recombinant protein to the core DRE element (CCGAC) provided evidence for a retarded band in the gel (lane 2), and for the absence of competition for binding using a test probe containing a mutated DRE element (lanes 7 and 8). When increasing concentrations (5 and 20 x) of test CRT probe for competition with DREB2D were applied, a gradual decrease was observed in the signal intensity of the retarded band (lanes 5 and 6), suggesting the second nucleotide position does not play a major role in protein-DNA interactions and that DREB2D is able to bind both DRE (T**A**CCGAC) and CRT (T**G**CCGAC) elements. DNA affinity purification sequencing (DAP-seq) assays were then conducted to identify the target genes of DREB2D and to refine the putative motifs bound by DREB2D. Significant DREB2D-binding peaks (fold-enrichment >3 and FDR<0.05) common to the three DAP-seq replicates were found in the 2 kb promoter region of 3,421 genes (Supplementary Table S14). This is illustrated in Fig. 7B with Cc05g12390 which encodes TRANVIA. GO-enrichment analysis performed on DREB2D target genes revealed significant enrichement in galactose metabolic processes, cellular glucose homeostasis and glucose binding (Supplementary Table S14). Moreover, many genes of the GMB coexpression network (32.2%) were identified as DREB2D target genes by DAP-seq analysis (Fig. 4, Supplementary Table S5). These genes are involved in all the different processes of CWP biosynthesis, including sucrose cleavage (SUSY), nucleotide sugar metabolism and transport (*GMD2*, *UAFT*), galactomannan synthesis (*MSR*, *GMGT*), cellulose synthesis (*CESA1*, *CESA3*, *CESA6*, *KOR1*), pectin synthesis and modification (*GAUT9*, *PME36*, *PMEI36*), secretory vesicle trafficking (*TVA*, *SCAMP4*, *RAB5a*, *RABD2a*) and cell wall protein synthesis (*AGP18*, *FLA17*). MEME analysis of 2 kb promoter sequences of the DREB2D target genes resulting from DAP-seq analysis identified two highly significant (p = 6E-113 and 1E-15, respectively) C-repeat motifs potentially bound by DREB2D, hereafter referred as to DREB2D-BS1 and –BS2 (Fig. 7C and D). Neither motif contained a distinct CCGAC core motif of DRE *cis*-acting elements. However, the second motif is very similar to that bound by AtDREB2D (Fig. 7D), with significant Tomtom motif-motif similarity (FDR=1.75E-06). The elements were then searched for in the proximal upstream region (2 kb) of eight selected genes of the GMB network representative of the different processes of CWP synthesis. All eight promoters contained either DREB2D-BS1 or DREB2D–BS2, or both motifs (Fig. 7C and D). For instance, DREB2D-BS1 was retrieved in the proximal upstream region of *UG4E1*, but DREB2D-BS2 was not, *GMGT1* was the opposite, as both motifs were identified at the same locations in the promoter of *MANS1*.

**Figure 7.**
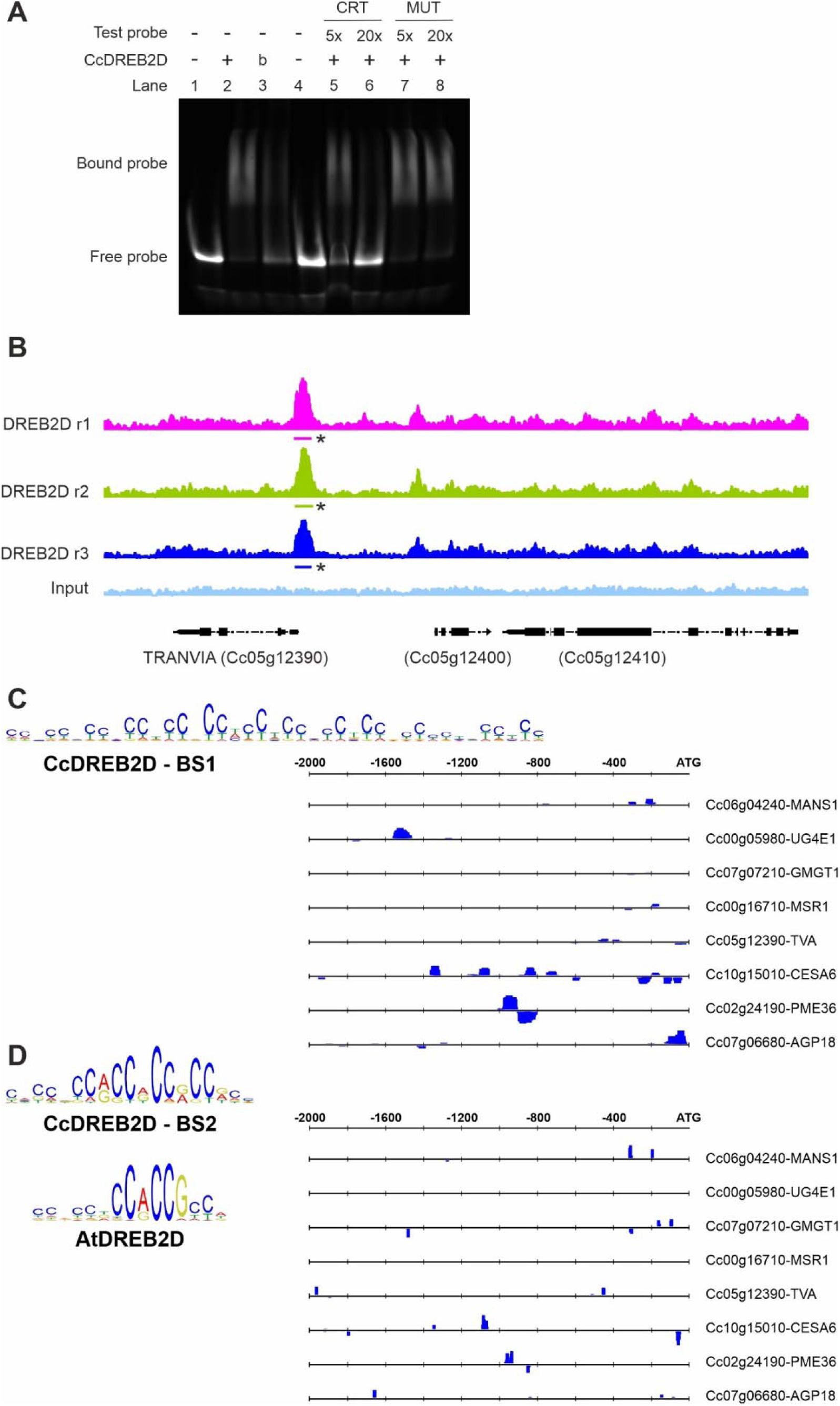
DREB2D binds the promoter of genes of the GMB coexpression network. **A.** Electrophoretic Mobility Shift Assay (EMSA) of DREB2D and dsDNA probes. Imaging was done by detection of the labelled DRE probe. CcDREB2D protein was omitted in lanes indicated by ‘-’, and included in lanes indicated by ‘+’, in lanes indicated by ‘b’ it was boiled for 15 min at 98 °C. Labelled DRE probe was present in each lane (200ng). Test probes (CRT, MUT) added to challenge the binding of the labelled DRE probe are indicated, (5x and 25x: five or 25 times excess compared to labelled DRE, ‘-’ indicates no added test probe). **B.** DAP-seq DNA-binding profile of DREBD2D (three replicates, r1-3) and the control (input). Horizontal bars below the plots represent the position of the significant peak regions in the promoter of *TRANVIA*. **C.** Logos of the two motifs (CcDREBD2D-BS1 and –BS2) significantly enriched in the promoter sequences of the DREBD2D target genes resulting from DAP-seq analysis. Number, position and strand of the two motifs in the proximal upstream region of eight selected genes of the GMB network.

## DISCUSSION

### Transcriptional orchestration of sucrose import and cleavage, nucleotide sugar metabolism, and synthesis, secretion and post-depositional modifications of galactomannans

Using a multi-location trial with a single *C. arabica* variety, and therefore environment as a source of variation in gene expression, we previously showed that five genes encoding enzymes of the core galactomannan biosynthetic machinery (MANS, MGT, GMGT, AGAL, UG4E) are quantitatively coexpressed across environments during seed development (Joët *et al*., 2014). To gain new insights into galactomannan biosynthesis, this finding prompted us to build the first gene coexpression network of the developing coffee endosperm. We chose to use the interspecific genetic variability present within the genus *Coffea* to obtain large variation in gene expression levels. Three results of the present study justified using this approach. First we showed that galactomannans and oil were the main seed storage compounds in all the coffee species studied. Second, considerable variation in the endosperm CWP content was observed among the fourteen coffee species, suggesting variation in the expression of genes involved in galactomannan accumulation, a condition required to build a coexpression network. Third, coffee species conserved a similar transcriptional endosperm development programme, despite the fact they started to diverge from each other between 4 and 8 million years BP (Bawin *et al*., 2021) and evolved towards great differences in seed development time.

The advantage of the mixed strategy (Guerin *et al*., 2016) used here to build a gene coexpression network, compared with a strict guide-gene approach (Joët *et al*., 2014), is being able to reveal novel key players of a biosynthetic pathway without prior information. Our gene coexpression network analysis first identified MSR1 in the module that contains the three guide genes MANS1, MGT1, and GMGT1. Gene coexpression network analysis also provided evidence for concerted transcriptional regulation, along with galactomannan biosynthesis, of sucrose import and cleavage activities (SWEET3, SUSY1), as well as of glycolytic activity (HXK-5, PGLYM-p), nucleotide sugar metabolism for the synthesis of GDP-man and UDP-gal (PPM, MGT1, UG4E1, UG4E2, UG4E3), and their import into Golgi (URGT6).

In addition to the upstream transcriptional control of central carbon metabolism and nucleotide sugar synthesis and transport which provide galactomannan building blocks, gene coexpression network analysis also revealed the tight orchestration of the multiple players involved in the functioning of Golgi and the trans-Golgi network, as well as the post-Golgi secretory pathway to the extracellular space. For instance, the GMB network included SCAMP4, one of the secretory carrier membrane proteins used as conventional markers for secretory vesicles, of which SCAMP2 is involved in the vesicular trafficking of pectin (Toyooka *et al*., 2009), plus key proteins STELLO and TRANVIA which play major roles in the assembly and delivery of CSCs. Similarly, RAB-GTPases, of which several members are present in the GMB network, play specialised roles in the transport of the different types of polysaccharides (cellulose, hemicellulose, pectin) from Golgi to the cell surface (Lunn *et al*., 2013). Furthermore, the GMB network includes the ECA4 component of the TPLATE adaptor complex, a major module for clathrin-mediated endocytosis (Gadeyne *et al*., 2014), for which evidence of a direct role in the recognition of cellulose synthase complexes for their internalization was recently put forward (Sánchez-Rodríguez *et al*., 2018). This is important since glycosyltransferases such as MANS and GMGT, require retrograde transport from secretory vesicles and late Golgi cisternae (trans) to earlier cisternae (cis) to maintain their steady-state spatial distribution across the Golgi stack (Zabotina *et al*., 2021). Finally, the GMB network revealed numerous receptor-like protein kinases which may help regulate CWP biosynthesis, since several receptor-like kinases have been shown to specifically control cellulose synthesis (Hématy *et al*., 2007) or to mitigate secondary cell wall thickening (Huang *et al*., 2018).

Coexpression analysis also highlighted the concerted regulation of galactomannan synthesis and post-depositional modifications, including the degree of galactose substitution (AGAL1), mannan remodelling (MAN2 and MAN6) through transglycosylation (Prakash *et al*., 2012) and acetylation (TBL38). Within the TBL family (DUF 231), TBL38 does not belong to subgroup II, whose members, including AXY4, are mannan O-acetyltransferases (MOATs) in Arabidopsis (Zhong *et al*., 2018). Rather, TBL38 belongs to subgroup IX, which includes PMR5, a functional acetyltransferase that mediates pectin acetylation (Chiniquy *et al*., 2019). The GMB network also includes three GDSL esterase genes close to the rice DARX1 protein, a Golgi-located enzyme that deacetylates the side chain of arabinoxylans and dynamically participates in acetylation/deacetylation of cell wall polymers during development (Zhang *et al*., 2017, 2019), acetylation of xylans being critical for interaction with cellulose fibrils in cell walls (Grantham *et al*., 2017).

### Intertwining of the transcriptional control exerted on the specific biosynthetic machineries of the different cell wall polysaccharides

The pathway-guided gene coexpression network revealed tight transcriptional coordination of the galactomannan biosynthesis machinery with those of the other components of coffee endosperm CWP, *i.e*. arabinogalactan proteins, cellulose, and, to a lesser extent, pectic polysaccharides (Redgwell and Fischer, 2006; Li *et al*., 2021). Two different sets of CESA complexes are mobilised for primary cell wall synthesis during cell expansion and secondary cell wall thickening, respectively (McFarlane *et al*., 2014). It is worth noting that the CESA genes which are connected to galactomannan biosynthetic genes, CESA1, CESA3, and CESA6-like genes (CESA2 and CESA6), are those encoding the three subunits that constitute the CSC recruited for the synthesis of the primary cell wall (Desprez *et al*., 2007; Persson *et al*., 2007). Assuming that lignin mainly occurs in the secondary cell wall, this important finding is in agreement with the absence of genes coding for phenylpropanoid biosynthesis and lignin polymerizing enzymes (laccases and peroxidases) in the GMB network, except peroxidase PRX12, which is thought to be involved in other processes than lignin biosynthesis in Arabidopsis (Hoffmann *et al*., 2020). Moreover, the GMB network contains several genes involved in the metabolism of pectin, which mainly accumulates in primary walls. Collectively, these results thus suggest that storage of polysaccharides in the coffee endosperm only takes place in primary walls. The results corroborate previous immunohistological observations of coffee endosperm cell walls showing that the β-1,4-mannan-specific monoclonal antibody BGM C6 labels cross the entire wall cross section (Sutherland *et al*., 2004). This contrasts with observations made in Arabidopsis where mannan epitopes were primarily located in the thickened secondary cell walls of xylem elements, xylem parenchyma and interfascicular fibres (Handford *et al*., 2003).

While the biosynthetic machinery of the different cell wall polysaccharides is highly coordinated at the transcriptional level, how coffee endosperm cells control the synthesis of large amounts of galactomannans in comparison with cellulose and pectin remains to be determined. The recruitment of three distinct UG4E genes (*UG4E1*, *UG4E2*, *UG4E3*) may help shift partitioning of most of the pool of UDP-Glu resulting from SUSY activity towards the synthesis of the galactomannan building block UDP-gal rather than cellulose, particularly in a cellular context where fatty acid synthesis also requires large supplies of sugar. In Arabidopsis, different coexpressed UG4E genes act synergistically for CWP synthesis (Seifert *et al*., 2002; Rösti *et al*., 2007). Moreover, transporters of specific nucleotide sugars in the Golgi apparatus usually work unidirectionnally, and are key rate-limiting control points towards different types of CWP (Kleczkowski and Igamberdiev, 2020), as shown with UDP-gal transporters (Rautengarten *et al*., 2014; Abedi *et al*., 2016).

### Mobilisation of an original set of master TFs for coffee endosperm maturation

LEC1-ABI3/FUS3/LEC2 (LAFL) proteins are well-characterised key regulators of embryo maturation, including oil and protein accumulation, in exalbuminous seeds (Fatihi *et al*., 2016; Alizadeh *et al*., 2021). Among these four so-called master regulators of seed development, only FUS3 was retrieved in the GMB network of the coffee endosperm and the functions of its coexpression partners in the network suggest it plays a central role in oil and CWP accumulation. It should be noted that there is no orthologue of AtLEC2 in the coffee genome. The role of LAFL regulators in seeds with persistent endosperm like coffee is very poorly documented (Miray *et al*., 2021). The function of FUS3 and ABI3 relatives has been suggested to be conserved in the dead endosperm of cereal caryopsis, but not that of LEC2 (Grimault *et al*., 2015). For instance, FUS3 displayed a conserved functionality in the barley endosperm for the control of storage protein accumulation, as shown by complementation of the loss-of-function *fus3* mutant in Arabidopsis (Moreno-Risueno *et al*., 2008). FUS3 is significantly expressed in the transient endosperm of Arabidopsis seeds, which also stores oil, though ten-fold less than that stored in the embryo (Barthole *et al*., 2014). In Arabidopsis, FUS3 controls the synthesis of storage lipids, either by directly activating fatty acid biosynthetic genes or indirectly through the activation of WRI1 (Wang and Perry, 2013).

WRI1 appeared to be a key regulatory hub in the GMB network of the coffee endosperm. WRI1 directly activates genes involved in late glycolysis and fatty acid (FA) synthesis during Arabidopsis seed maturation (Baud *et al*., 2007). WRI1 is thought to play a ubiquitous role in oil synthesis in plants (Ma *et al*., 2013). In the present work, we observed that numerous coexpression partners of WRI1 are involved in fatty acid synthesis, glycolysis and in the TCA cycle, suggesting it may play a similar role in the coffee endosperm. Little is known about the role of WRI1 in seeds with persistent endosperm. Transcript profiling showed it is expressed concomitantly with fatty acid biosynthetic genes in the oily endosperm of oil palm (Dussert *et al*., 2013), *Paeonia ostia* (Xiu *et al*., 2018) and castor bean (Yang *et al*., 2019). The ability of the oil palm endosperm-specific WRI1-2 to trigger FA synthesis has already been validated using transient expression in tobacco leaves (Dussert *et al*., 2013) and castor WRI1 shown to complement Arabidopsis *wri1* mutants (Yang *et al*., 2019).

Our results thus suggest a conserved role for FUS3 and WRI1 as master TFs for storage compound accumulation in the cellular endosperm of coffee seeds. It is worth noting that WRI1 not only functions downstream of LEC2 (Baud *et al*., 2007) but also of FUS3 in Arabidopsis (Yamamoto *et al*., 2010). In addition to the canonical role of FUS3 and WRI1, the massive orientation of the carbon flow towards galactomannans may have been assisted by the selection of a particular assemblage of TFs during plant evolution. The identification of SHN2 in the GMB network is particularly striking in this context. Indeed, members of the SHINE clade of AP2 domain transcription factors were first shown to activate wax biosynthesis and to alter cuticle properties in Arabidopsis (Aharoni *et al*., 2004). However, they were later also described as key regulators of cellulose and hemicellulose biosynthesis pathways in rice and tobacco (Ambavaram *et al*., 2011; Liu *et al*., 2017). When ectopically expressed in these species, Arabidopsis and poplar SHN2 were shown to coordinate the up-regulation of cellulose and other primary cell wall compound biosynthesis, as well as the down-regulation of lignin biosynthesis and many secondary NAC and MYB TFs that trigger secondary wall biosynthesis (Zhong and Ye, 2015). Microscopy and chemical analyses of sclerenchyma cells of rice leaves overexpressing AtSHN2 further revealed a marked increase in cell wall thickness (around 45%) compared to the wild type, mainly due to the increased deposition of cellulose and hemicelluloses in place of lignin (Ambavaram *et al*., 2011). Given that the gene regulatory network associated with SHN2, with both activator and repressor functions, appears to be highly conserved in both monocot and dicot plants, and that the present coffee galactomannan-guided co-expression network mimics large parts of this transcriptional repertoire, it is tempting to attribute a similar role to SHN2 in triggering the peculiar primary cell wall thickening programme of coffee endosperm parenchyma cells, in particular through the recruitment of secondary TFs specialised in the activation of galactomannan biosynthetic genes.

The timing of cell wall thickening also presumably relies on precise specialised crosstalk between phytohormones and their related transcriptional networks which determine the division and expansion of coffee endosperm parenchyma cells. In this respect, the detection of ARF4 and ARF8 in the GMB network suggests that auxin is involved in triggering the onset of endosperm filling. ARFs have been extensively reported to play a key regulatory role in fruit and seed development (Cao et al., 2020). In Arabidopsis, ARF2 and -8 are transcription factors that inhibit cell division and seed growth. Deletion mutations in AtARF2 cause Arabidopsis seeds to become abnormally large (Schruff *et al*., 2006). Compared with the WT, the arf8 mutant had longer/larger capsules and more seeds, suggesting that seed size and capsule form are regulated by ARFs in Arabidopsis (Goetz *et al*., 2006). ARF4 plays a role in determining tomato fruit cell wall architecture (Sagar *et al*., 2013). What is more, ARFs function not only in the auxin signalling pathway but also in many other phytohormone signal transduction pathways in plants, including ABA, gibberellins and brassinosteroids (Cancé *et al*., 2022). For instance, AtARF2 is required for the normal brassinosteroid response and the brassinosteroid-regulated BRASSINOSTEROID INSENSITIVE 2 (BIN2) kinase can increase the expression of auxin-induced genes by directly inactivating AtARF2 (Vert *et al*., 2008). The presence of this key brassinosteroid-associated gene in the GMB network is therefore of interest.

### DREB2D, a secondary TF that plays a critical role in rewiring nucleotide sugar metabolism to galactomannan precursor synthesis

The present study provides considerable experimental evidence that DREB2D is a key TF for cell wall synthesis in the coffee endosperm: (i) it is the only TF of module 2 of the GMB network that groups numerous genes of the core galactomannan biosynthetic machinery, (ii) DREB2D is the only TF that is a direct partner of the guide genes MANS1, GMGT1, and MGT1, (iii) somatic embryos overexpressing DREB2D display significant up-regulation of numerous key genes of galactomannan biosynthesis and modification, including those encoding UG4E1, UG4E2, UG4E3, AGAL1, MSR1, TRANVIA, PPM, TBL38, TBL42 and mannan endo-1,4-beta-mannosidases, (iv) DAP-seq analysis of DREB2D target genes identified many genes of the GMB coexpression network involved in sucrose cleavage, nucleotide sugar metabolism and transport, galactomannan synthesis, cellulose synthesis, pectin synthesis and modification, secretory vesicle trafficking and cell wall protein synthesis.

However, the three key genes MGT1, MANS1 and GMGT1 were not up-regulated in DREB2D+ somatic embryos. This suggests that additional cellular/molecular factors are required to fully activate the galactomannan biosynthetic machinery and trigger their accumulation. In this respect, the functional interaction between the TFs HaDREB2 and HaHSFA9 is worth mentioning, as it synergistically trans-activates target sHSP genes during sunflower seed development (Díaz-Martín *et al*., 2005). When overexpressed in tobacco, HaDREB2 alone does not up-regulate sHSP genes, whereas, when co-expressed with HaHSFA9, it increases sHSP accumulation and improves seed longevity (Almoguera *et al*., 2009). Interestingly, HSFA9 was identified in our ST3-ST5 coexpression network and may be an interesting candidate for further investigation of galactomannan biosynthesis in coffee.

In the absence of functional galactomannan machinery in somatic embryos overexpressing DREB2D, the dramatic up-regulation of UG4E genes presumably led to increased production of UDP-galactose and its diversion to other pathways such as the biosynthesis of raffinose family oligosaccharides. Indeed, raffinose and stachyose contents were found to be significantly increased in DREB2D+ lines. These results corroborate our previous findings in *C. arabica* seeds in which raffinose and stachyose were identified as a transient storage form that buffers excess UDP-galactose before it is remobilised and supplies auxiliary sources of building blocks for galactomannan synthesis (Joët et al., 2014). Since GolS1, which catalyses the first reaction of the raffinose family oligosaccharide pathway, uses UDP-gal and *myo*-inositol as a substrate, this could be a simple explanation for the smaller amounts of *myo*-inositol detected in DREB2D+ lines. Our results therefore point to a major role for DREB2D in controlling UDP-gal homeostasis during CWP biosynthesis in coffee.

## Abbreviations

DAP-Seq: DNA affinity purification sequencing
GMB: galactomannan biosynthesis.

## SUPPLEMENTARY DATA

**Table S1.** RNAseq statistics of samples used for gene coexpression analysis.

**Table S2.** List of primers used for qRT-PCR.

**Table S3.** EMSA probe sequences, showing the DREB target sites in bold, mutations in bold italics.

**Table S4.** Fatty acid composition of mature seeds.

**Table S5**. The 174 genes forming the galactomannan biosynthesis co-expression network.

**Table S6.** The 144 genes forming the galactomannan co-expression network built using ST3-ST5 RNA-seq data.

**Table S7.** Number of partners, at different R thresholds, of 10 key transcription factors of the GMB network.

**Table S8.** qPCR analysis of galactomannan-related genes in coffee somatic embryos of lines overexpressing DREB2D and in controls.

**Table S9.** RNAseq statistics of coffee in coffee somatic embryos overexpressing DREB2D and in controls.

**Table S10.** Normalized gene expression (rpkm) in coffee somatic overexpressing DREB2D and in controls.

**Table S11.** Genes significantly up-regulated (531) and down-regulated (350) in coffee somatic embryos overexpressing *DREB2D* tested using DESeq2 analysis.

**Table S12.** MAPMAN functional enrichment analysis of differentially expressed genes (DEGs) in lines overexpressing DREB2D+ lines compared to in controls.

**Table S13.** List of differentially expressed genes whose expression levels were correlated with that of the DREB2D transgene (R^2^ > 0.9).

**Table S14.** List of the 3,589 significant DREB2D-binding peaks shared by the three DAP-seq replicates found in the 2 kb promoter region of 3,421 genes.

**Table S15.** GO-enrichment analysis of the 3421 DREB2D-target genes.

**Supplementary Figure S1.** Multivariate analysis of the transcriptomes of the 69 combinations of species × developmental stage. A. Hierarchical cluster analysis. B. Score plots of Principal Components PC1 versus PC2 of the principal component analysis (PCA).

**Supplementary Figure S2.** Stages 3-5 of the galactomannan-guided gene coexpression network. The network was constructed using the 42 transcriptomes (ST3-ST5 seed developmental stages), six galactomannan-related guide genes (*MANS1*, *GMGT1*, *MGT1*, *UG4E1*, *UG4E2*, *UG4E3*), two rounds of coexpression analysis and an |R| threshold of 0.85. Each of the 144 genes of the network was assigned to a functional group, for which the most relevant ones are shown in colour.

**Supplementary Figure S3.** Expression profiles (rpkm) across seed development of key seed maturation transcription factors (TFs) in the 14 *Coffea* species studied.

**Supplementary Figure S4.** Somatic embryos regenerated from lines overexpressing DREB2D and in controls.

**Supplementary Figure S5.** Phylogenetic analysis of DREB2 *Arabidopsis thaliana* and *Coffea canephora* genes. The phylogenetic tree was constructed using the amino acid sequence of Cc10g02270 and Arabidopsis DREB transcription factor (TF) orthologues using the Plaza platform (HOM04M00166 family)(Van Bel *et al*., 2022). Multiple sequence alignment using MUSCLE and PhyloXML tree construction algorithm as described in Kreft *et al*. (2017).

## ACKNOWLEDGMENTS

The authors thank the *Coffea* Biological Resources Center (BRC Coffea, maintained by IRD and CIRAD in Reunion Island) for providing the plant material and the Plant Protection Platform (3P, IBiSA) for hosting visiting researchers. The authors are also grateful to Dr. Sylvain Santoni and to the staff of the AGAP Institut platform for preparing the cDNA libraries. The authors acknowledge the ISO 9001 certified IRD i-Trop HPC (member of the South Green Platform) at IRD Montpellier for providing HPC resources that have contributed to the research results reported within this paper.

## AUTHOR CONTRIBUTIONS

TJ, SD: conceptualization; AKS, JS, VV, MCC, HE, SR, PL, TJ, SD: methodology; AKS, FM, SR, PL, TJ, SD: formal analysis; AKS, JS, VV, MCC, EL, TJ: investigation; TJ, SD: resources; AKS, TJ, SD, PL, FM: data curation; TJ, SD, AKS : writing - original draft; JS, VV, MCC, FM, EL, SR, HE, PL: writing - review & editing; TJ, SD: visualization; TJ, SD: supervision. All the authors approved the final version of the manuscript.

## CONFLICT OF INTEREST

The authors declare they have no conflict of interest.

## FUNDING

AKS was financially supported by the European Molecular Biology Organisation (EMBO) under a long-term fellowship grant [ALTF 478-2017]. MGX acknowledges financial support from France Génomique National infrastructure, funded as part of “Investissement d’Avenir” program managed by Agence Nationale pour la Recherche [ANR-10-INBS-09].

## Notes

### Competing Interest Statement

The authors have declared no competing interest.

